# Insights into standards of care – dexamethasone and antibodies against COVID-19 in hamster models

**DOI:** 10.1101/2021.12.17.473180

**Authors:** Emanuel Wyler, Julia M. Adler, Kathrin Eschke, Gustavo Teixeira Alves, Stefan Peidli, Fabian Pott, Julia Kazmierski, Laura Michalick, Olivia Kershaw, Judith Bushe, Sandro Andreotti, Peter Pennitz, Azza Abdelgawad, Dylan Postmus, Christine Goffinet, Jakob Kreye, S Momsen Reincke, Harald Prüss, Nils Blüthgen, Achim D. Gruber, Wolfgang M. Kuebler, Martin Witzenrath, Markus Landthaler, Geraldine Nouailles, Jakob Trimpert

**Affiliations:** Berlin Institute for Medical Systems Biology (BIMSB), Max Delbrück Center for Molecular Medicine in the Helmholtz Association (MDC), Berlin, Germany; Institute of Virology, Freie Universität Berlin, Berlin, Germany; Charité - Universitätsmedizin Berlin, Corporate Member of Freie Universität Berlin and Humboldt-Universität zu Berlin, Institute of Pathology, Berlin, Germany; IRI Life Sciences, Institute for Biology, Humboldt-Universität zu Berlin, Berlin, Germany; Charité - Universitätsmedizin Berlin, Corporate Member of Freie Universität Berlin and Humboldt-Universität zu Berlin, Institute of Virology, Berlin, Germany; Berlin Institute of Health (BIH), Berlin, Germany; Charité - Universitätsmedizin Berlin, Corporate Member of Freie Universität Berlin and Humboldt-Universität zu Berlin, Institute of Physiology, Berlin, Germany; Institute of Veterinary Pathology, Freie Universität Berlin, Berlin, Germany; Bioinformatics Solution Center, Freie Universität Berlin, Berlin, Germany; Charité - Universitätsmedizin Berlin, Corporate Member of Freie Universität Berlin and Humboldt-Universität zu Berlin, Division of Pulmonary Inflammation, Berlin, Germany; Charité - Universitätsmedizin Berlin, Corporate Member of Freie Universität Berlin and Humboldt-Universität zu Berlin, Department of Infectious Diseases and Respiratory Medicine, Berlin, Germany; German Center for Neurodegenerative Diseases (DZNE) Berlin. Helmholtz Innovation Lab BaoBab (Brain Antibody-omics and B-cell Lab); Charité - Universitätsmedizin Berlin, Corporate Member of Freie Universität Berlin and Humboldt-Universität zu Berlin, Department of Neurology and Experimental Neurology, Berlin, Germany; Charité - Universitätsmedizin Berlin, Corporate Member of Freie Universität Berlin and Humboldt-Universität zu Berlin, Department of Pediatric Neurology, Berlin, Germany

**Keywords:** COVID-19 treatment, dexamethasone, antibody, monoclonal antibody therapy

## Abstract

**Rationale:** In face of the ongoing SARS-CoV-2 pandemic, effective and well-understood treatment options are still scarce. While vaccines have proven instrumental in fighting SARS-CoV-2, their efficacy is challenged by vaccine hesitancy, novel variants and short-lasting immunity. Therefore, understanding and optimization of therapeutic options remains essential.

**Objectives:** We aimed at generating a deeper understanding on how currently used drugs, specifically dexamethasone and anti-SARS-CoV-2 antibodies, affect SARS-CoV-2 infection and host responses. Possible synergistic effects of both substances are investigated to evaluate combinatorial treatments.

**Methods:** By using two COVID-19 hamster models, pulmonary immune responses were analyzed to characterize effects of treatment with either dexamethasone, anti-SARS-CoV-2 spike monoclonal antibody or a combination of both. scRNA sequencing was employed to reveal transcriptional response to treatment on a single cell level.

**Measurements and main results:** Dexamethasone treatment resulted in similar or increased viral loads compared to controls. Anti-SARS-CoV-2 antibody treatment alone or combined with dexamethasone successfully reduced pulmonary viral burden. Dexamethasone exhibited strong anti-inflammatory effects and prevented fulminant disease in a severe COVID-19-like disease model. Combination therapy showed additive benefits with both anti-viral and anti-inflammatory potency. Bulk and single-cell transcriptomic analyses confirmed dampened inflammatory cell recruitment into lungs upon dexamethasone treatment and identified a candidate subpopulation of neutrophils specifically responsive to dexamethasone.

**Conclusions:** Our analyses i) confirm the anti-inflammatory properties and indicate possible modes of action for dexamethasone, ii) validate anti-viral effects of anti-SARS-CoV-2 antibody treatment, and iii) reveal synergistic effects of a combination therapy and can thus inform more effective COVID-19 therapies.

## Introduction

A novel coronavirus (CoV), severe acute respiratory distress syndrome CoV-2 (SARS-CoV-2) emerged in December 2019 in Wuhan, China and evolved rapidly into an ongoing pandemic (1). While development of vaccines was successful, there is still a lack of approved, effective and well-understood CoV disease 2019 (COVID-19) treatments (2, 3).

To devise successful host-directed therapeutic strategies, understanding of COVID-19 pathogenesis is required. For COVID-19 patients, virus-triggered exuberant cytokine release and associated tissue damage play a crucial role in disease severity, e. g. elevated levels of pro-inflammatory cytokines as well as loss of effector T-cells were associated with fatal outcomes (4–7). Despite growing knowledge regarding the mechanisms of severe disease, very few treatment options are available, so that the use of corticosteroids, specifically dexamethasone, remains the treatment of choice for many critically ill patients.

Initially, use of corticosteroids was not recommended in treatment guidelines due to their broadly immunosuppressive action (8–10). Evidently, glucocorticoid treatment can result in impaired virus clearance (11). Nevertheless, in the RECOVERY trial, clinical application of dexamethasone yielded positive effects, especially for COVID-19 patients requiring oxygen therapy (12). Although corticosteroids are now used routinely to treat critically-ill COVID-19 patients, putative hazards for mild to moderate COVID-19 patients as well as mechanisms underlying its protective efficacy in severe COVID-19 remain obscure and only begin to be investigated in greater depth (13).

Since the development of small molecule inhibitors of virus replication is all but trivial, passive immunization using monoclonal antibodies (mAb) became an important approach to COVID-19 therapy relatively early in the pandemic. SARS-CoV-2 cell entry inhibition by mAb targeting the receptor-binding domain (RBD) of the spike protein revealed high effectivity (14). Various anti-SARS-CoV-2 antibodies have been developed and are currently tested in *in-vivo* models or in clinical trials (15–17). The first approved anti-SARS-CoV-2 mAb is REGN-COV2 a combination of the mAbs casirivimab and imdevimab. Effectivity depends on timing of therapy, as application early in disease can prevent patient hospitalization. Early therapy or prophylaxis reduces virus titers in the respiratory tract and consequently the risk of severe disease progression (18, 19). The therapeutic activity of mAbs depends critically on the presence of their binding sites in currently circulating virus variants (20).

Dexamethasone, in contrast, acts non-specifically on the hosts’ immune response and is less likely to lose therapeutic power to new variants if induced immune responses remain similarly pathogenic. To date, detailed understanding of the mechanisms behind the action of these two standard treatments is still not fully developed. To fill this knowledge gap, we examined the therapeutic effects of dexamethasone and monoclonal anti-SARS-CoV-2 antibody treatment as well as their potential as synergistic combinatorial therapy in hamster models of moderate and severe COVID-19 using single-cell and bulk transcriptome-based analyses.

## Methods

An online Supplement is provided giving more details on the here described methods.

### Ethics statement and COVID-19 hamster models

Experiments including female and male hamsters (*Mesocricetus auratus*; breed RjHan:AURA, JanvierLabs, France) and Roborovski hamster (*Phodopus roborovskii*, via German pet trade) were approved and executed in compliance with all applicable regulations (permit number 0086/20). SARS-CoV-2 preparation (21) and infection of hamsters were carried out as previously described (22, 23). Treatments were applied as single i.p. treatment with 30 mg/kg mAb as previously described (16) and daily i.m. treatment with 2 mg/kg dexamethasone in the respective groups. Hamsters were monitored daily until they reached scheduled take-out time points or defined humane endpoints. Virus titers and RNA copies were determined by plaque assay and quantitative RT-PCR analysis as previously described (22).

### Histopathology and *in situ*-hybridization of SARS-CoV-2 RNA

For histopathology and *in situ*-hybridization (ISH), lungs were processed, and tissues evaluated by board-certified veterinary pathologists in a blinded fashion following standardized recommendations, including pneumonia-specific scoring parameters as described previously (24).

### Annotations of the *M. auratus* and *P. roborowskii* genome

The *M. auratus* genome was derived from Ensembl and modified as previously described (25). The detailed description of the *de-novo* gene assembly of the Roborovski hamster genome was deposited on a pre-print server (26).

### Bulk RNA analysis

For RNA-Bulk Sequencing of both hamster species, the right medial lung lobe was removed and RNA isolated using Trizol reagent according to the manufacturer’s instructions. Bulk RNA sequencing libraries were constructed using the Nebnext Ultra II Directional RNA Library Prep Kit (New England Biolabs) and sequenced on a Nextseq 500 or Novaseq 6000 device. Reads were aligned to the genome using hisat2 (27) and gene expression quantified using quasR (27).

### Single-cell-RNA-Sequencing

To enable scRNA-Seq, cells were isolated from Roborovski hamsters’ caudal lung lobe as previously described (25). 1,000,000 lung cells per sample were subjected to CMO labeling according to manufacturers’ instructions (3’-CellPlex-Kit-Set-A, 10x Genomics). Labelled cells from 12 samples were pooled, filtered and counted. Pooled cells were adjusted to a final concentration of ~1,600 cells / μL, and 197,760 cells were split into four equal pools and subjected to partitioning into Gel-Beads-in-Emulsions with the aim of recovering a maximum of 120,000 single cells from four lanes by following the instructions of Chromium-Next-GEM-Single-Cell-3’-Reagent-Kits-v3.1 (Dual Index) provided by the manufacturer (10x Genomics). Library sequencing was performed on a Novaseq 6000 device (Illumina), with SP4 flow cells (read1: 28 nucleotides, read2: 150 nucleotides). Sequencing of one of four libraries failed.

### Analysis of single-cell-RNA-Sequencing data

Analysis of the single-cell data was based on Seurat (28). Raw and processed data is available through GEO at https://www.ncbi.nlm.nih.gov/geo/query/acc.cgi?acc=GSE191080, code through Github at https://github.com/Berlin-Hamster-Single-Cell-Consortium/Dwarf-Hamster-Dexamethasone-Antibody. Details on single cell analysis and RNA velocity analysis can be found in the online Supplement.

## Results

### Dexamethasone treatment prevents severe disease, while monoclonal antibodies decrease viral burden

Following SARS-CoV-2 infection, Syrian hamsters lost body weight. Irrespective of treatment, Syrian hamsters failed to show significant differences in body weight development nor did they present with severe signs of disease (Figure 1A, B). Titers of replication competent virus of all hamsters receiving mAb or combination treatment was below the detectable level at all sampling time points. The use of dexamethasone alone significantly increased viral titers in the lungs of Syrian hamsters and delayed viral clearance with moderately increased titers on day 5 and sustained increased titers 7 days post infection (dpi) (Figure 1C). This effect was also evident in virus gRNA levels in the lungs (Figure 1D), but not in the upper respiratory tract (Figure 1E), which is the common site of sampling in patients.

**Figure 1.**
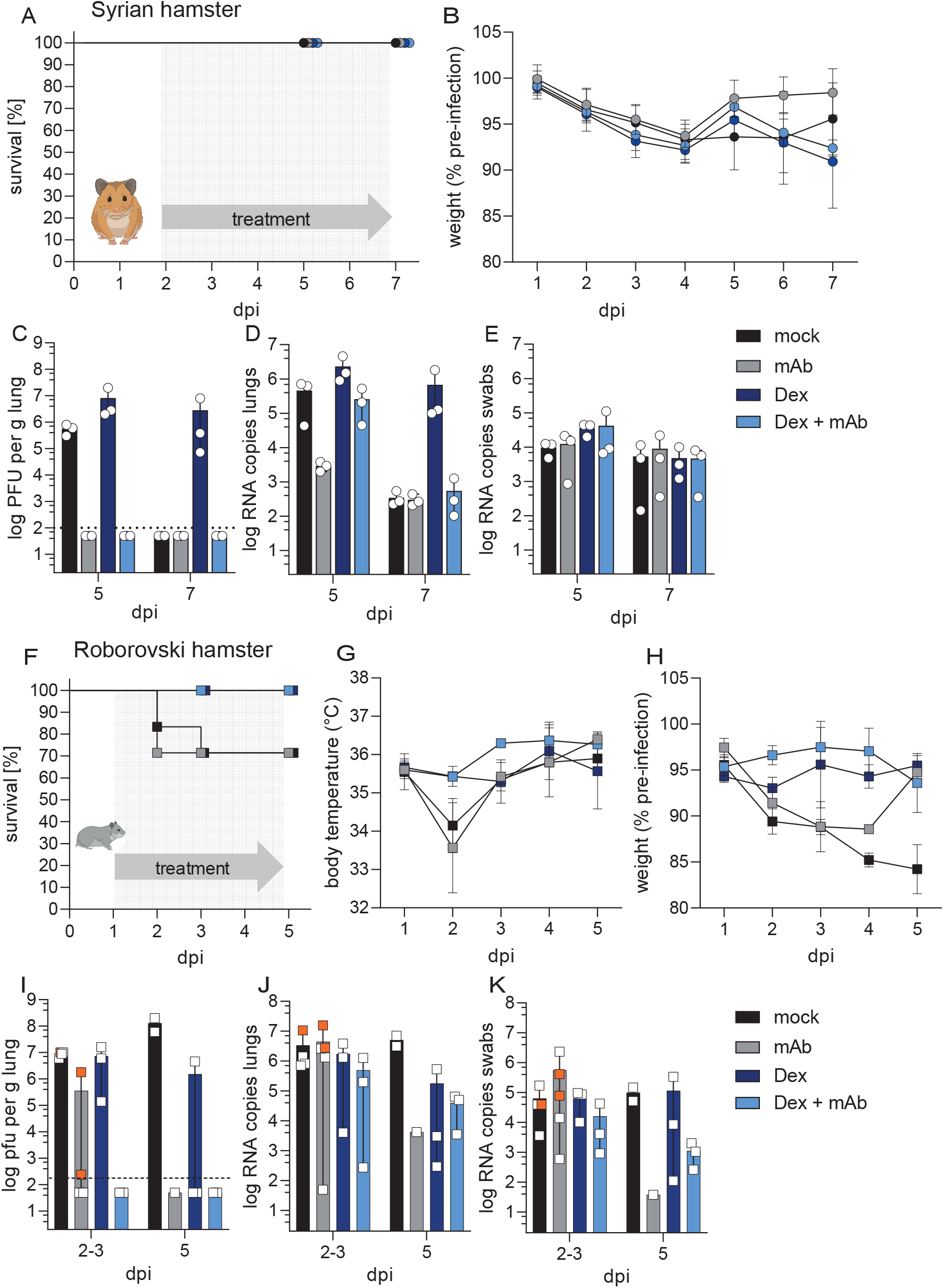
Clinics of SARS-COV-2 infected Syrian and Roborovski hamsters under COVID-19 therapy. Syrian hamsters (**A – E**) were challenged with SARS-CoV-2 (1 x 10^5^ pfu Wildtype (WT)) and treated once at 2 dpi with 30 mg/kg mAb CV07-209 (mAb, n = 6), daily starting at 2 dpi with 2mg/kg Dexamethasone (Dex, n = 6) or received combination treatment (Dex + mAb, n = 6). Survival rates (**A**) in percent of SARS-CoV-2 infected Syrian hamsters and body weight (**B**) development in percent after virus challenge were measured until analysis time point (5 dpi, n = 3 and 7 dpi n = 3) and displayed according to treatment group. (**B**) Results are displayed as mean ± SD. (**C**) Quantification of replication competent virus as plaque-forming units (pfu) per gram homogenized lung tissue. Dotted line marks the limit of detection (DL = 100 pfu). Titers below the detection limits were set to DL/2 = 50 pfu. Number of genomic RNA (gRNA) copies detected in homogenized lung tissue (**D**) and oropharyngeal swabs (**E**). (**C – E**) Results are shown as mean with range. Roborovski hamsters (**F – K**) were challenged with SARS-CoV-2 (1 x 10^5^ pfu Wildtype (WT)) and treated once at 1 dpi with 30 mg/kg mAb CV07-209 (mAb, n = 6), daily starting at 1 dpi with 2 mg/kg Dexamethasone (Dex, n = 6) or received combination treatment (Dex + mAb, n = 6). Survival rates (**F**) in percent of SARS-CoV-2 infected Roborovski hamsters, body temperature (**G**) in degree Celsius and body weight (**H**) development in percent after virus challenge were measured until planned analysis time point (3 dpi, and 5 dpi) or until termination due to score sheet criteria (non-survivors) according to treatment group. Two hamsters from the mAb group and one hamsters from the mock-treated group were euthanized at 2 dpi (represented by orange squares (**I – K**)). One hamster from the mock-treated group reached end point criteria at 3 dpi and was included in 3 dpi time point analysis as planned. (**G**, **H**) Results are displayed as mean ± SD. (**I**) Virus titers displayed as pfu per gram homogenized lung tissue. Dotted line marks the limit of detection (DL = 100 pfu). Titers below the detection limits were set to DL/2 = 50 pfu. J + K) Quantification of gRNA copies in homogenized lung tissue (**J**) and oropharyngeal swabs (**K**). (**I – K**) Results are displayed as mean with range.

Contrary to Syrian hamsters, Roborovski hamsters which develop fulminant disease early after infection (23) displayed marked differences in clinical parameters in response to specific treatments. Specifically, both dexamethasone alone and in combination with mAb protected Roborovski hamsters from severe disease progression. By contrast, hamsters assigned to mAb treatment (2/6 on 2 dpi) and animals receiving mock treatment (2/6 on 2 dpi or 3 dpi) had to be euthanized prior to the terminal time point as they reached human endpoint criteria (Figure 1F). Hamsters that developed severe disease in respective groups presented with drastic drops in body temperature at 2 dpi (Figure 1G). Until the end of the experiment, body weights in the dexamethasone treated groups remained stable, animals in the mAb treatment group recovered from initial weight losses, while mock treated animals continued to lose weight throughout the experiment (Figure 1H). Similar to Syrian hamsters, the viral load in the lungs of Roborovski hamsters treated with either mAb or combinatorial therapy was below the detectable level at all regular sampling days. Only Roborovski hamsters that had to be terminated at 2 dpi showed high titers of replication competent virus despite mAb treatment (Figure 1I). In contrast to the results obtained from Syrian hamsters, no boost of viral replication was observed in the dexamethasone treated group of Roborovski hamsters compared to mock treated animals. This result was evident for all time-points, on both replicating virus and virus gRNA level in the lungs as well as in the upper respiratory tract (Figure 1J, K).

### Dexamethasone restricts the inflammatory response

Dexamethasone is a useful drug to treat severe COVID-19 patients (12). To better characterize effects on local pathomechanisms, we performed lung histopathology upon dexamethasone, mAb and combinatorial therapy against SARS-CoV-2 in models of moderate (Syrian hamster) and severe (Roborovski hamster) COVID-19 (Figure 2A – F).

**Figure 2.**
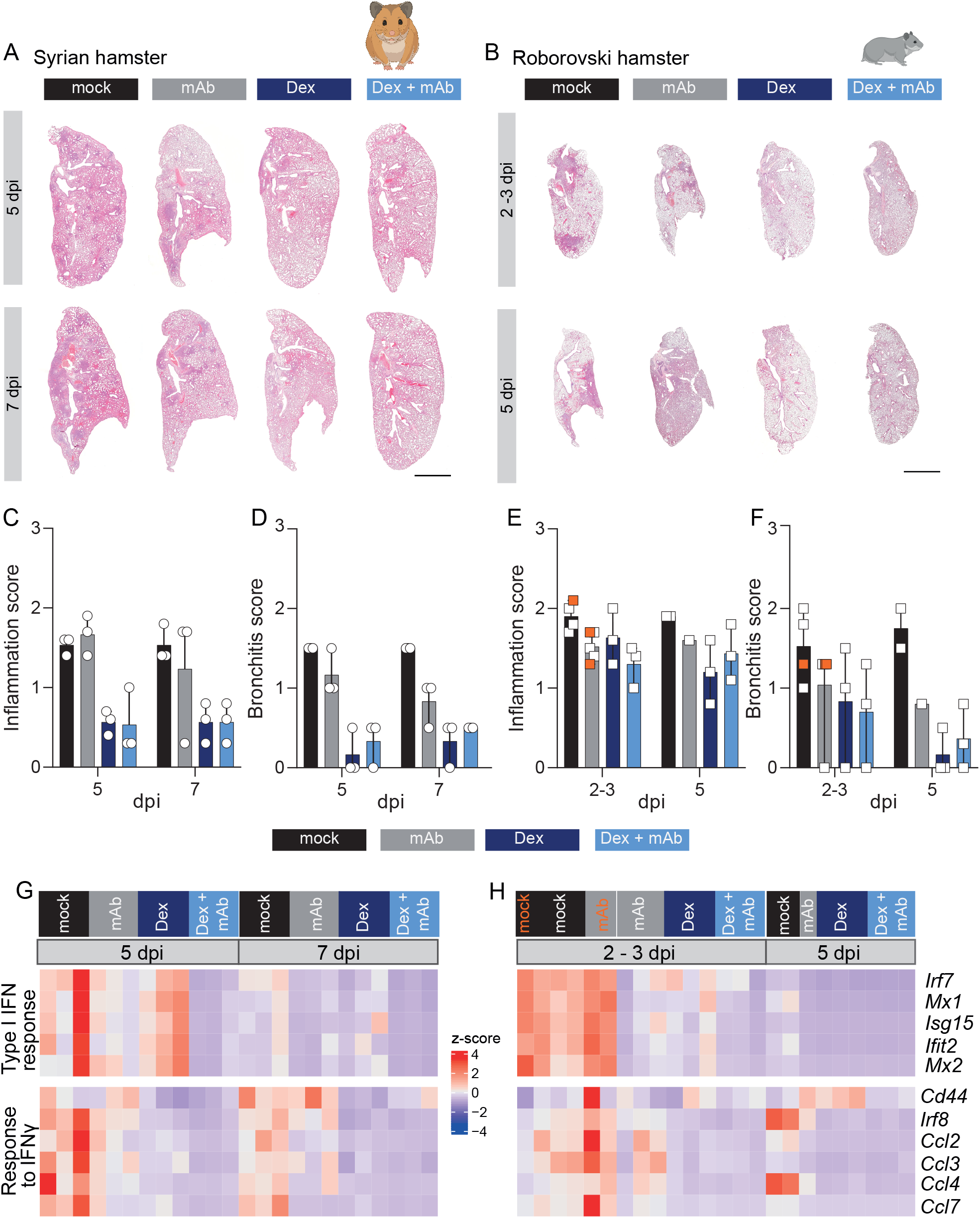
Dexamethasone treatment dampens inflammatory responses of SARS-CoV-2 infected hamsters. Longitudinal sections of H&E-stained left lungs from representative Syrian hamsters (**A**) and Roborovski hamsters (**B**) at indicated time points post infection. Consolidated areas indicative of pneumonia appear in darker colours. Bars = 3 mm. Lung inflammation score ((**C**) Syrian and (**E**) Roborovski hamsters) accounting for the severities of pneumonia, immune cell influx, perivascular lymphocyte cuffs, bronchitis, bronchial epithelial necrosis, alveolar epithelial necrosis and type II pneumocyte hyperplasia. Bronchitis score ((**D**) Syrian and (**F**) Roborovski hamsters) assessing bronchitis and bronchial epithelial necrosis. Gene expression ((**G**) Syrian and (**H**) Roborovski hamsters) was quantified using polyA RNA high-throughput sequencing from Syrian hamster lung samples. Shown are z-scores of fpkm (fragments per kilo base of transcript per million mapped fragments) values calculated over all samples on a colour scale ranging from blue (−4) to red (+4) for selected genes. Time points and treatments are shown on top of the heatmap. Samples from animals euthanized at 2 dpi are shown in orange.

Lung histology indicated that in both Syrian (Figure 2A) and Roborovski hamsters (Figure 2B) dexamethasone and combination treatment markedly reduced immune cell infiltrates over time (Figure E1). Inflammation and bronchitis scores were reduced at from 5 dpi on in all groups receiving dexamethasone, which corresponds to 3 or 4 days post treatment start for Syrian and Roborovski hamster, respectively (Figure 2C – F). mAb treatment alone reduced pneumonia, however, to a lesser extent as compared to dexamethasone (Figures 2A – F, E1).

Next, we investigated how anti-viral and inflammatory transcriptional responses were influenced by treatment in Syrian (Figure 2G, E2A) and Roborovski hamsters (Figure 2H, E2B) over time. Therefore, we analysed previously established viral infection related gene sets, *response to type I interferon* (*IFN*) and *interferon-gamma* (*IFN-γ*) *(25, 29)*. In Syrian hamsters, the amplitude of the *type I IFN response* genes decreased from 5 to 7 dpi in the absence of treatment (Figure 2G, E2A). mAb treatment alone or in combination with dexamethasone led to further reduction in gene expression of the *type I IFN response* genes. In contrast, *IFN-γ response* set genes decreased more upon dexamethasone compared to mAb treatment (Figure 2G, E2A). Similar effects were observed in Roborovski hamsters (Figure 2H, E2B). The combination treatment led to a strong reduction of both gene sets, independent of hamster species (Figure 2G,H).

Taken together, treatment related improvement in clinical parameters and histopathology correlated with substantially altered gene expression profiles in general, and a reduced expression of the *response to IFN-γ* gene set following dexamethasone treatment specifically.

### Dexamethasone reduces influx of immune cells and stabilizes endothelial cells

As described above, both mAb and dexamethasone treatment, and in particular their combination, attenuated inflammatory aspects of pneumonia following SARS-CoV-2 infection, thereby rescuing Roborovski hamsters from an otherwise fatal disease course.

In order to investigate cellular mechanisms underlying these treatment effects, we next performed pulmonary scRNA-seq of Roborovski hamsters for all treatment groups at 3 dpi. First, we evaluated the absolute content and composition of cell types by measuring total cell counts of the dissociated tissue (Figure 3A) and relative cell type distribution from scRNA-seq data (Figure 3B–D, E3A-J). Lungs from dexamethasone (alone or in combination with mAb) treated hamsters yielded significantly lower total cell counts (Figure 3A). This reduction likely originated from reduced infection-triggered pulmonary immune cell immigration. NK cell numbers were significantly lower in dexamethasone treated groups compared to mock and mAb treated hamsters; similarly, neutrophil, monocytic macrophage, *Treml4*^+^ monocyte, T and B cell showed reduced numbers in hamsters receiving dexamethasone, although the difference was not statistically significant (Figure 3B, C). Notably, endothelial cells had significantly higher counts in groups treated with a combination therapy of dexamethasone and mAb (Figure 3D) as compared to mock treated animals. Higher endothelial cell counts were likely caused by mechanisms governing endothelial protection, rather than cell proliferation, since increased *Mki67* and *Top2* expression was not detectable in endothelial cells (Figure E3K). The notion of endothelial protection was supported by histopathological analyses showing reduced edema formation and reduced endothelialitis in dexamethasone treated groups (Figure 3E, F upper panel), thus replicating findings in patients (30). Histopathological analyses likewise confirmed reduction of recruited immune cells following single dexamethasone treatment alone, and in combination with mAb (Figure 3F). In contrast to mAb treatment alone, dexamethasone therefore largely reduced recruitment of immune cells.

**Figure 3.**
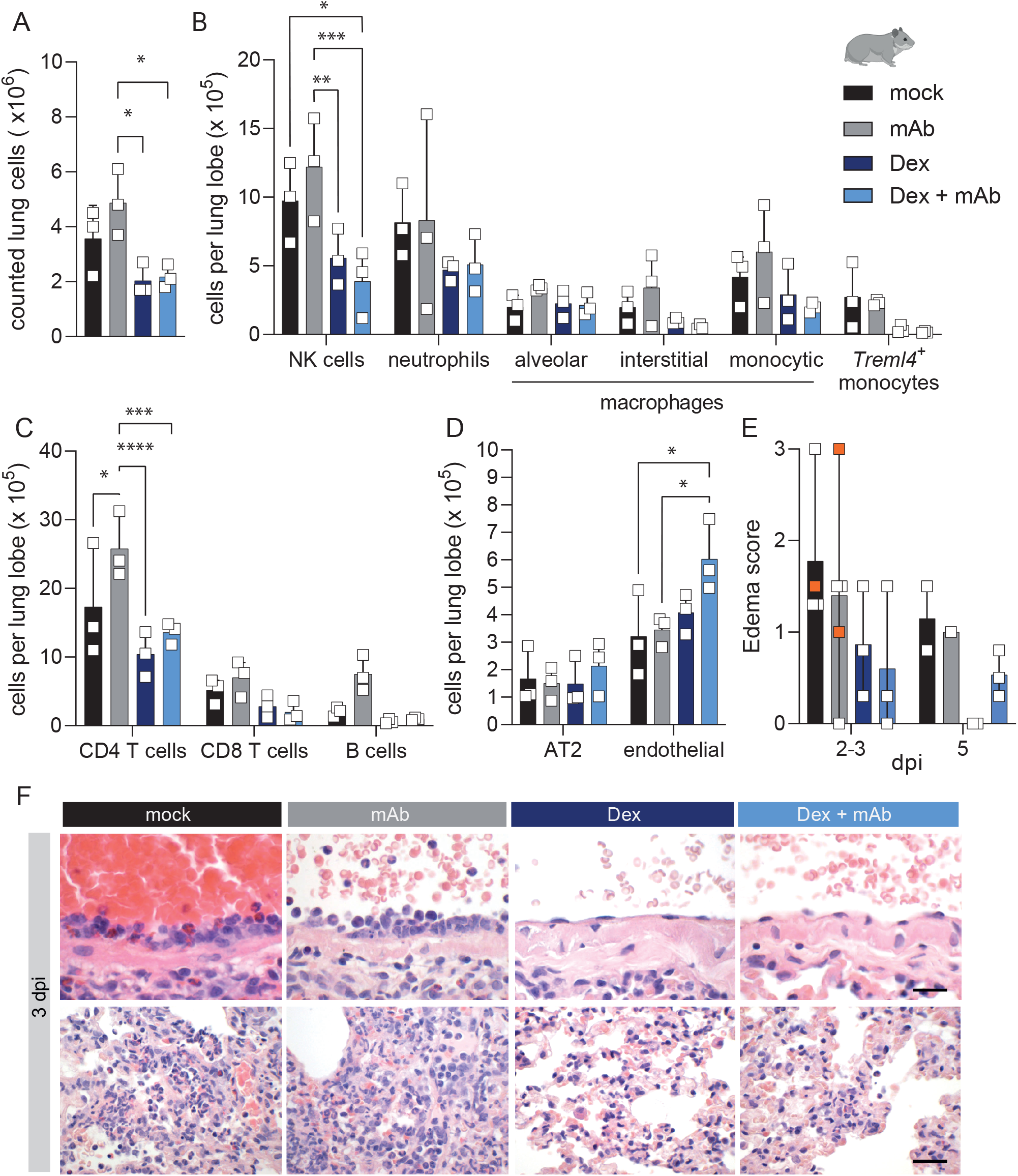
Dexamethasone limits immune cell recruitment in Roborovski hamsters. Roborovski hamsters were challenged with SARS-CoV-2 (1 x 10^5^ pfu Wildtype (WT)), treated once at 1 dpi with 30 mg/kg mAb CV07-209 (mAb), daily starting at 1 dpi with 2 mg/kg Dexamethasone (Dex) or received combination treatment (Dex + mAb). At 3 dpi n = 3 Roborovski hamsters of each group were subjected to pulmonary single-cell RNA sequencing analysis. Pulmonary single cell suspensions were generated, cells were microscopically counted and total numbers per lung lobe calculated. (**A**) Cell count of isolated cells per lung lobe according to treatment group. Calculated numbers of indicated innate immune cells (**B**), T and B lymphocytes (**C**) and AT2 and endothelial cells (**D**) based on scRNA-seq determined cell frequencies (Figure E3) and according to treatment group. Data display means ± SD. n = 3 per group. (A – D) Two-way ANOVA, Tukey’s multiple comparisons test. * P < 0.05, ** P < 0.01, *** P < 0.001, **** P < 0.0001. (**E**) Edema Score resulting from semi-quantitative assessment of alveolar and perivascular edema (**F**) H&E stained histopathology of pulmonary vascular endothelia (upper panel) and lung parenchyma (lower panel) from Roborovski hamsters at 3 dpi. Mock and mAb treated groups had moderate to marked endothelialitis with activation and loss of endothelial cells whereas the vascular endothelium remained mostly intact in Dex and Dex + mAb treated groups. The inflammatory response was more pronounced in mock and mAb treated hamsters compared to Dex and Dex + mAb treated animals. Differences were particularly observed for infiltrating neutrophils, macrophages and, lymphocytes as well as for the degree of alveolar epithelial cell necrosis. Size bars: 15 μm (top) and 25 μm (bottom).

### Neutrophils and monocytic macrophages exhibit strong responses to dexamethasone

Dexamethasone directly impairs transcription of NF-kB target genes via Rela/p65 and Crebbp/CBP (31). In order to assess the effect of dexamethasone treatment, known target genes of the glucocorticoid receptor, the *coagulation cascade factor F13a1* (32), the plasma apolipoprotein *serum amyloid a-3 protein (Saa3*) (33), and *Dusp1/MKP-1*, an inhibitor of the MAP kinase pathways (34), were investigated (Figure E4A-C). Neutrophils and macrophages, particularly monocytic macrophages, from dexamethasone treated groups, showed strong increase in target gene expression, *F13a1*, *Dusp1*, and *Saa3* (Figure E4A – C).

For an unbiased view of the data, we selected all genes that were at least four-fold upregulated in all cell types (Figure 4A). Again, monocytic macrophages and neutrophils stood out with several upregulated genes, including *Saa3* and *F13a1* as mentioned above. We identified a dexamethasone-induced transcriptional program common to several cell types, whereas some genes, for example *Gal* (coding for galanin and galanin message-associated peptides) in endothelial cells were cell-type specific. In contrast, tissue cells, including endothelial cells, alveolar epithelial cell type 2 (AT2) or smooth muscle cells did not show substantial upregulation of gene expression in response to dexamethasone alone (Figure 4A). Notably, the mRNA of the glucocorticoid receptor, encoded by the *Nr3c1* gene, is ubiquitously present in both Roborovski hamsters and Syrian hamsters, and not modulated by SARS-CoV-2 infection or the employed treatments (Figure E4D).

**Figure 4.**
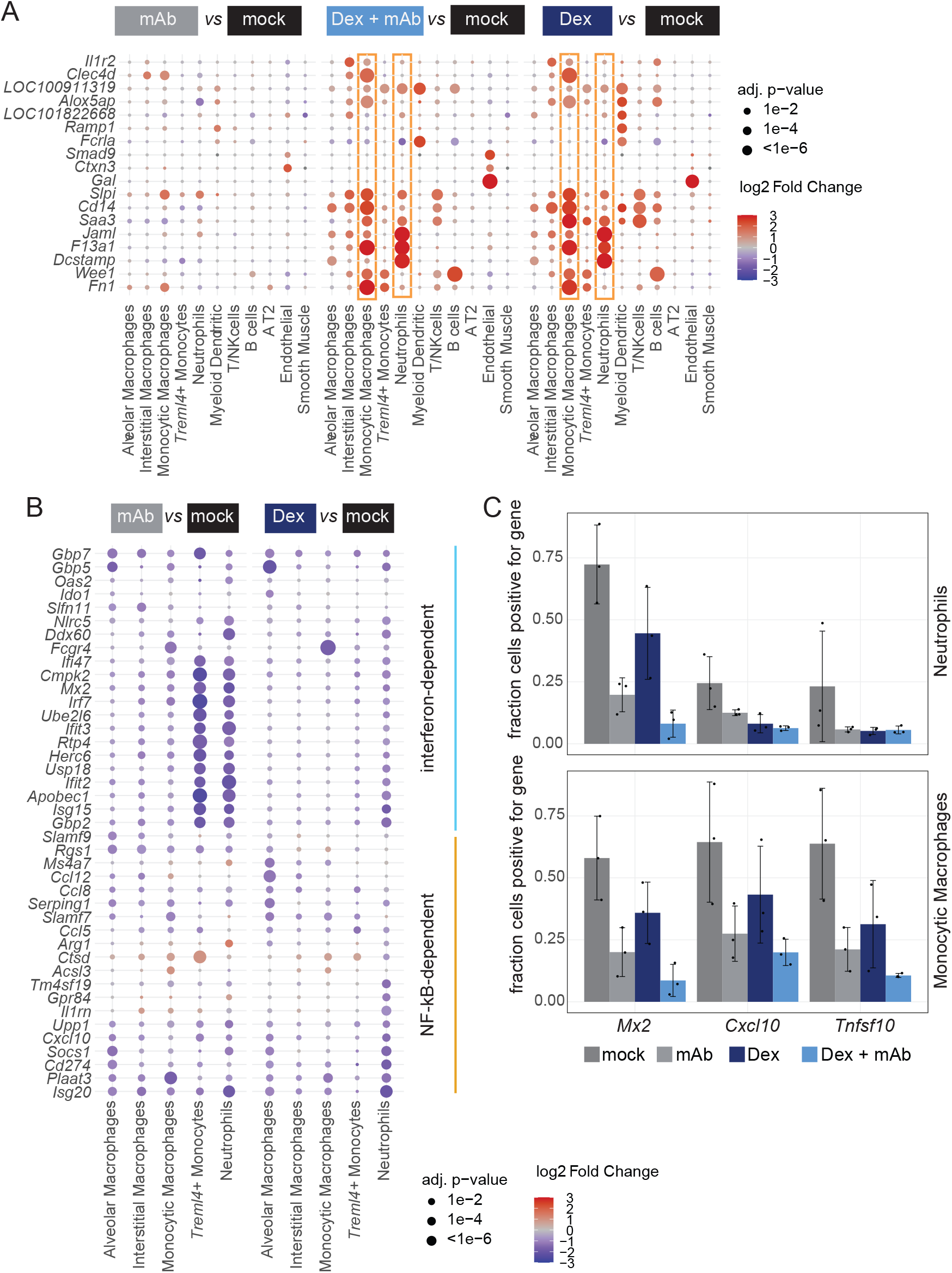
Macrophages and neutrophil show strongest gene expression changes by Dexamethasone treatments. (**A**) Shown are genes with at least four-fold upregulation in at least one cell type in the dexamethasone compared to the mock-treated animals, for all three treatments compared to mock. Size and colors of the dots indicate log2-transformed fold changes (FC) and p-values, respectively. Adjusted (adj) p-values were calculated by DEseq2 using Benjamini–Hochberg corrections of two-sided Wald test p-values. Genes are ordered by unsupervised clustering. (**B**) Shown are interferon and NF-kB-dependent genes as determined in Figure E4 for the comparisons Dex vs. mock and Dex + mAB vs. mock. Otherwise as in (**A**). (**C**) Expression of *Mx2*, *Tnfsf10*, and *Cxcl10* in neutrophils (top) and monocytic macrophages (bottom). Shown are the fraction of cells with ≥ one mRNA count (means ± SD. n = 3 per group).

Next, we asked which disease-relevant changes in gene expression were influenced by treatment in different cell types. We therefore assessed changes in gene expression between treatments for each cell type in an unbiased manner (Figure E4E). We noticed consistent downregulation of a group of interferon-induced genes (ISGs) such as *Ifit2/3*, *Ifi27*, *Ifi209* in animals treated with mAb alone or in combination with dexamethasone, but not with dexamethasone alone. Conversely, some genes, such as *Tnfsf10* (coding for the pro-inflammatory cytokine Trail) in neutrophils were more reduced in dexamethasone treated compared to mAb treated animals.

In order to understand the changes in gene expression patterns caused by these treatments, we defined, based on our Syrian hamster scRNA-seq data (25), two groups of gene sets. The first was viral PAMP dependent (identified as “NF-kB-dependent”), the second induced by the infection in general (“interferon-dependent”) (Figure E4F). Whereas the “interferon-dependent” gene expression was reduced more by mAb compared to dexamethasone treatment, for the “NF-kB-dependent” gene set we in tendency observed the opposite (Figure 4B). We scrutinized this effect in detail in monocytic macrophages and neutrophils, and found that in neutrophils, the NF-kB-driven cytokine genes *Cxcl10* and *Tnfsf10* are particularly decreased by dexamethasone, whereas the ISG *Mx2* was specifically diminished by mAb treatment (Figure 4C). For all genes, the combination treatment showed an additive effect (Figure 4).

Overall, these data suggest that the reduced viral load in mAb-treated animals leads to a generally reduced antiviral/type 1 interferon signal, whereas dexamethasone treatment downregulates specific genes in some cell types, such as the pro-inflammatory cytokines *Tnfsf10* and *Cxcl10* in neutrophils, thereby attenuating classic features of pneumonia in animals receiving dexamethasone.

### Dexamethasone alters the neutrophilic response to SARS-CoV-2 infection

Given that neutrophils are critical drivers of immune pathology and showed a particularly strong reactivity to dexamethasone treatment, we investigated this cell type in greater detail. For this, we sub-clustered the neutrophil population into 11 subpopulations (Figure 5A).

**Figure 5.**
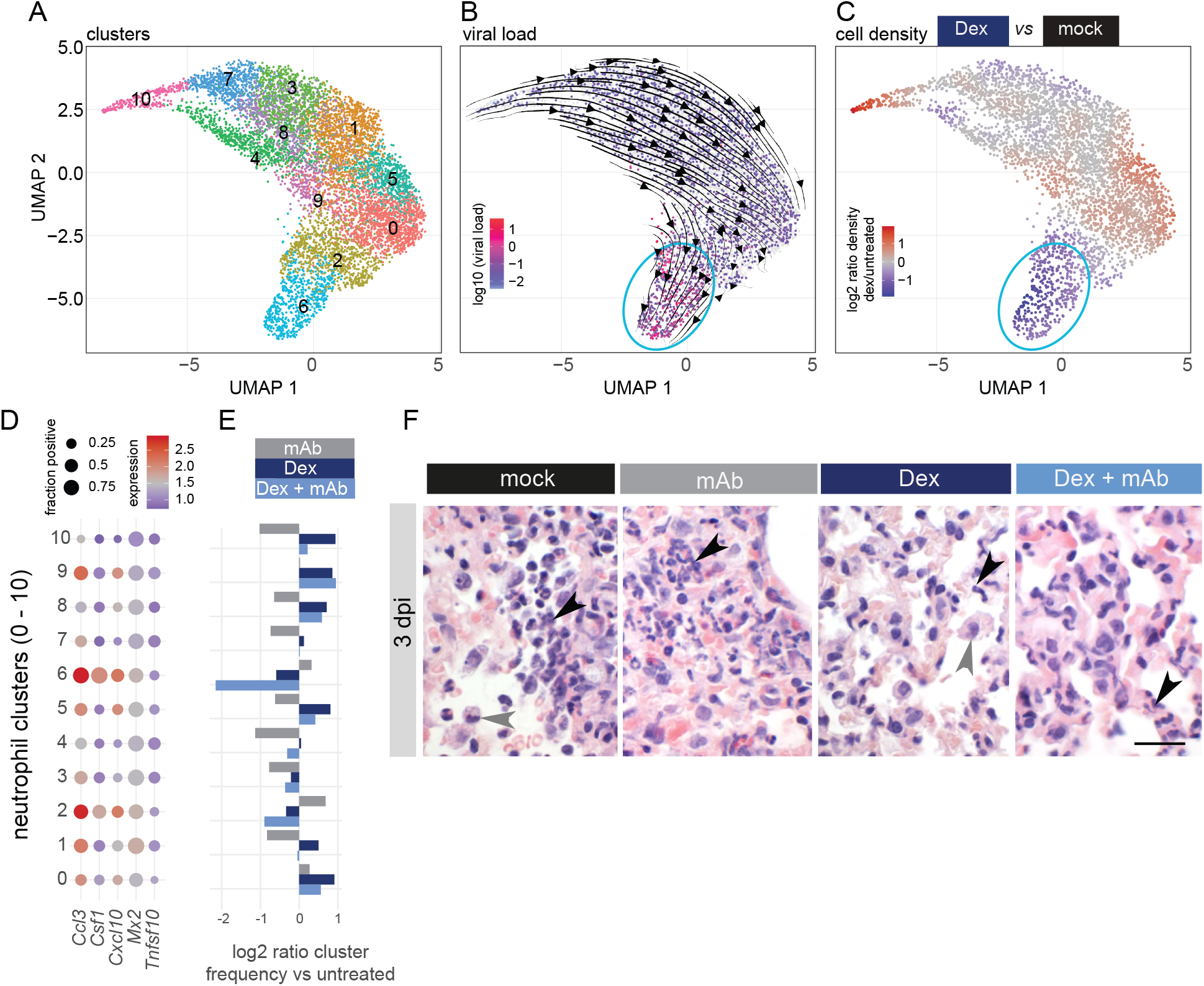
Absence of a specific chemokine-expressing subset of neutrophils upon dexamethasone treatment in Roborovski hamsters. (**A**) Neutrophils from the scRNA-seq data were sub-clustered using the Louvain algorithm based on their individual transcriptomes, and two-dimensional projections performed using the UMAP algorithm. Cells were coloured by their cluster identity. (**B**) Projection as in (**A**), but cells are coloured by the log10-transformed percentage of viral RNA. Overlaid are the stream arrows derived from the RNA velocity analysis. Neutrophil cluster 6 is marked with a light blue oval. (**C**) Changes in cellular density on the UMAP projection were calculated, and cells coloured by fold changes of the indicated Dex vs. mock. Red indicates increased density, and blue indicates decreased density. Neutrophil cluster 6 is marked with a light blue oval. (**D**) Graph indicates the log2-transformed fold changes of the cell counts in the respective neutrophil clusters 1-10, with all three treatments compared to mock. For example, in cluster 6 there are about one third less cells (dark blue bar at −0.6, which corresponds to log2 of 0.66) upon dexamethasone treatment. (**E**) Dot plots show the expression of selected genes over all hamsters in the clusters as defined in (**A**). The dot size indicates the fraction of cells in the clusters as indicated on the left from mock-treated animals, with ≥ one mRNA count for the respective gene. The colour represents the average expression in those cells. (**F**) Histopathology of Roborovski hamsters 3 days after infection revealed moderate to marked alveolar and interstitial infiltration with viable and degenerate neutrophils (black arrowheads) in mock and mAb treated animals as well as elevated numbers of alveolar macrophages (gray arrowhead). Dex and Dex + mAb treated hamsters had lower numbers of neutrophils especially in their alveolar spaces and mild to moderate numbers of neutrophils in alveolar capillaries (black arrowheads). Activated alveolar macrophages phagocytized cellular debris and cleared the inflammatory response (gray arrowhead). Scale bar = 20 μm.

In order to understand the transcriptional dynamics within neutrophils and the influence of the treatments used here, we performed an RNA velocity analysis which can predict the future state of individual cells (35, 36). This showed a transcriptional trend towards the cluster on the bottom of the projection (cluster 6 in Figure 5A), which also showed a particularly high viral RNA content (Figure 5B, E5A). Importantly, cell density in that cluster decreased upon dexamethasone treatment (Figure 5C, E5B).

Among the genes that were particularly prominent in cluster 6 were the cytokines and macrophage and lymphocyte attractants *Csf1* and *Cc13* (37, 38) (Figure E5C). We therefore plotted the expression of these two genes along with the ISG/NF-kB targets *Mx2/Tnfsf10/Cxcl10*, which showed that neutrophils in cluster 6 express *Csf1* and *Ccl3* at particularly high levels (Figure 5D), in the same time, these cells become less abundant upon dexamethasone and particularly combination treatment (Figure 5E). Concomitantly, by histopathology analysis, we observed less neutrophils in the dexamethasone treated groups (Figure 5F). Of note, cells expressing mRNAs of receptors (*Csf1r*, *Ccr1*, *Ccr4*, and *Ccr5*) corresponding to cytokines *Csf1* and *Ccl3* were less abundant in the lungs upon dexamethasone treatment (Figure E5D, compare with Figure 3B and E3B). In addition, neutrophil-cluster 6 showed particularly low and high expression of *Il1r2* and *Isg20* (Figure E5E), respectively, thereby recapitulating the phenotypes seen for immunosuppressive and IFN^active^ neutrophils in the peripheral blood of COVID-19 patients (13).

To generalize the observation of this transcriptional dynamic, we applied diffusion map analysis of neutrophils to identify their most prominent direction of variation (39, 40) (Figure E5F). For each treatment, we show the neutrophil density along the diffusion axis (Figure E5G, upper part), which we defined as the first non-trivial component of the diffusion map. The directional progression towards the right on this axis (which is the same cellular state represented as neutrophil-cluster 6 above) is present in all conditions, as shown by the average RNA velocity projected onto the diffusion axis (Figure E5G, lower part). However, most neutrophils derived from hamsters treated with dexamethasone or combinatorial treatment are found at the leftmost part of the axis, whereas neutrophils from hamsters with mAb and mock treatment are split into a left and right part, confirming that with dexamethasone treatment an otherwise directional progression of neutrophils is limited. In order to relate the diffusion axis to biological effects, we scored hallmark signatures (41) for every neutrophil and linearly correlated each hallmark with the diffusion axis (Figure E5H, upper part). In addition, we correlated the expression profiles of each gene with the diffusion axis (Figure E5H, lower part). These correlations revealed that the drive towards neutrophil-cluster 6 marked by high expression of *Csf1* and *Ccl3* and elevated amounts of viral RNA is accompanied by an increase of interferon/inflammatory response gene expression (such as *Isg15* or *Cd274*), and a decrease in the levels of classical neutrophil marker genes such as *S100a8/9* or *Pglyrp1*. Dexamethasone limits this dynamic, effectively keeping the neutrophils in a stationary transcriptomic state at the left part of the diffusion axis. As we will discuss in detail, this stagnation could be a reason for the reduced production of lymphocyte attractants and consequently, the reduction of lung infiltrates.

## Discussion

In this study, we examined the effects of separate and combined anti-viral and anti-inflammatory treatments for COVID-19 in two hamster models reflecting a moderate (Syrian hamster) and severe (Roborovski hamster) disease course, respectively. Using histopathology and bulk and single-cell transcriptomic analysis of hamsters subjected to dexamethasone, monoclonal antibody, and combination treatment, we demonstrate treatment efficacy, and identified a subset of neutrophils that express macrophage/lymphocyte attracting cytokines and can be impeded by dexamethasone. The use of dexamethasone caused a boost of virus replication and a significant delay of viral clearance in Syrian hamsters, albeit without significantly worsening the clinical course of disease. In the light of existing literature on the enhanced replication of respiratory viruses upon dexamethasone treatment (42), and data that overall shows a tendency towards a boost of SARS-CoV-2 replication in dexamethasone treated patients (11, 43–45), this result is not unexpected and may imply a risk for increased and/or prolonged transmissibility. Still, dexamethasone exerted the expected anti-inflammatory effects and attenuated inflammatory lung injury. As previously reported (16), the mAb CV07-209 employed in this study effectively abolished virus replication within 48 hours of treatment. At the dose applied here, the mAb inhibited the boost of virus replication after dexamethasone treatment. This suggests that a combination of dexamethasone and mAb may present an effective way to reduce inflammation and at the same time suppress virus replication, limiting the risk of viral transmission. This would advocate for the use of a combination therapy in patients at risk of severe disease relatively early when active virus replication is still ongoing, and before lung injury or COVID-19 triggered fibrosis (46) develop. Interestingly, the use of dexamethasone in the Roborovski hamster, a species highly susceptible to severe COVID-19-like disease, did not boost virus replication at any of the examined time points. One possible explanation could be that the virus-restrictive immunity targeted by dexamethasone in Syrian hamsters is dysregulated in Roborovski hamsters, and consequently its inhibition has no impact on viral control.

Treatment of SARS-CoV-2 infected hamsters with dexamethasone reduced the extent of lung infiltrates, comparable to what can be observed in CT-scans of human COVID-19 patients (47). In the single-cell RNA-seq analysis, this effect was evident as reduced abundance of infiltrating leukocytes and lymphocytes. In an unbiased comparison of gene expression patterns in the different lung cell types, we found that neutrophils are particularly affected by dexamethasone treatment. A detailed analysis showed that upon SARS-CoV-2 infection, neutrophils move towards a state with high expression of the cytokines *Csf1* and *Ccl3*, and that this movement is impaired by dexamethasone. Furthermore, the receptors of the two cytokines are expressed on a range of cell types that become less abundant in the lungs upon dexamethasone treatment. This together suggests a mechanistic link underlying the protective effect, through reduction of lung infiltrates, by dexamethasone. These results are in line with the key role of neutrophils in COVID-19 pathogenesis (48), and corroborate recent findings highlighting the effect of dexamethasone on neutrophils in peripheral blood (13). Although neutrophils in blood and lung might not be directly comparable, the observation by Sinha and colleagues, a neutrophil “IFNactive” program restrained by dexamethasone, was similarly observed in the present study.

In addition to its effects on PMN, dexamethasone treatment exerted protective effects on the endothelium of SARS-CoV-2 infected hamsters, likely by reducing endothelial injury caused by cytotoxic immunity and bystander effects conveyed by the pro-inflammatory program executed by highly stimulated immune cells. As a secondary effect, the expression of inflammatory mediators by endothelial cells could also be reduced. Of clinical relevance, endothelial protection will reduce the development of lung edema and micro-thrombosis and may thus contribute to improved gas-exchange in dexamethasone treated patients.

Care should be taken not to transfer findings from animals uncritically to patients. Yet, it should be noted that we and others recently demonstrated comparability between immunological responses and pulmonary phenotypes in hamsters and humans in response to SARS-CoV-2 infection (25, 49, 50). That notwithstanding, future studies should ideally compare patient data to the findings reported here with the obvious constraint of limitations in the availability of corresponding human biomaterial.

In summary, we found that broadly active anti-inflammatory and immunosuppressive agents such as dexamethasone may have a strong benefit in SARS-CoV-2 infection at high risk for severe disease when applied before the onset of severe illness, particularly when combined with an anti-viral agent. A recent analysis showed that COVID-19-related ARDS patients can be classified into hypo- and hyperinflammatory types, with corticosteroid treatment being beneficial only for the latter (51). Animal models as the ones described here can help to better dissect causes and types of COVID-19 lung pathologies, and thus help to improve therapeutic strategies.

## Acknowledgement

Computation has been performed on the HPC for Research cluster of the Berlin Institute of Health and the max cluster of the Max Delbruck Center. The authors thank Angela Linke, Michaela Scholz, and Simon Dökel for excellent technical assistance with histopathology and ISH, and Jeannine Wilde, Madlen Sohn, and Tatiana Borodina (MDC Scientific Genomics Platforms) for sequencing. Hamster icons were used from BioRender.com.

