## Supplemental Methods and Figures for "Insights into standards of care – dexamethasone and antibodies against COVID-19 in hamster models"

*Virus*

Prior to infection, SARS-CoV-2 isolate (BetaCoV/Munich/BavPat1/2020) grown on Vero E6 cells in minimal essential medium (MEM; PAN Biotech) with 10% fetal bovine serum (PAN Biotech) and 100IU/ml Penicillin G and 100 µg/ml Streptomycin (Carl Roth). After incubation and titre determination, stocks were prepared and stored at -80°C. Integrity of the furin cleavage site was confirmed through sequencing of stocks prior to infection.

*Infection*

To ensure successful intranasal infection, hamsters were anaesthetised with a triple anaesthesia consisting of midazolam (2 mg/kg), butorphanol (2.5 mg/kg) and medetomidine (0.15 mg/kg). Subsequently $1x10^5$ PFU SARS-CoV-2 diluted in 60 µl MEM (PAN, Biotech) for Syrian hamsters (1) or 30 µl for Roborovski hamsters (2) were applied intranasally.

*Animal Experiment*

The hamsters were randomly assigned to 4 groups. Treatment started on day 2 after infection for Syrian hamsters and day 1 after infection for Roborovski hamsters. Group 1 received placebo therapy, whilst group 2 was treated with a single dose of anti-SARS-CoV-2 antibody CV-07-209 (2) injected intra-peritoneally at a dose of 30 mg/kg. Hamsters of the third group were treated daily with dexamethasone (2 mg/kg applied intra-muscularly. In group 4 the animals received a combination of both therapies described for group 2 and 3.

Hamsters were weighted daily and monitored for signs of disease twice-daily. Severely sick animals were euthanized according to defined humane endpoints including body temperature <33 °C, acute respiratory distress or weight loss >20%. Hamsters were selected randomly for all scheduled take-out time points (day 3 and 5 for Roborovski hamsters, day 5 and 7 for Syrian hamsters). Take out timepoints were scheduled according to the different disease courses on day 5 and 7 for Syrian hamsters and on day 3 and 5 for Roborovski dwarf hamsters. For euthanasia, animals were anesthetised with medetomidine (0.15 mg/kg), midazolam (2 mg/kg), and butorphanol (2.5 mg/kg) prior to exsanguination (1). To conduct virological, histopathological and single-cell sequencing analysis, serum, EDTA blood, lungs and oropharyngeal swabs were collected.

*RNA extraction and qPCR*

RNA was extracted from oropharyngeal swabs and 25 mg homogenized lung tissue. To do so innuPREP Virus DNA/RNA Kit (Analytic Jena, Jena, Germany) was used according to manufacturer's instructions. For quantification of viral RNA, we used NEB Luna Universal Probe One-Step RT-qPCR Kit (New England Biolabs) and the qPCR was conducted on the StepOnePlus Real-Time PCR System (Thermo Fisher Scientific) (3).

*Plaque assay*

Titration was performed to quantify replication competent virus. Briefly, lung tissue was homogenized in a bead mill (Analytic Jena). Thereafter, 50 mg of homogenized lung were used to prepare 10-fold dilutions starting from - 1 to - 6 and transferred onto Vero E6 cells seeded in 24-well plates (Sarstedt, Nümbrecht, Germany). Subsequently, the plates were incubated for 2,5 hours at 37°C and overlayed with 1,5% methylcellulose (Sigma Aldrich). After 72 hours cells were fixed in 4% formaldehyde, stained with 0.75% crystal violet and results were evaluated by counting the plaques.

*Analysis of single-cell-RNA-Sequencing data*

Analysis of the single-cell data was based on Seurat (4). All used code with annotation is available through Github at https://github.com/Berlin-Hamster-Single-Cell-Consortium/Dwarf-Hamster-Dexamethasone-Antibody.

Details on single cell analysis and RNA velocity analysis can be found below:

Briefly, samples were integrated following SCtransform (5). Cell types were annotated based on the expression of marker genes in Louvain clusters. Variability and statistical tests were calculated based on three animals per group. Within the neutrophil subclustering, differential analysis was based on Louvain clusters.

For RNA velocity analysis, sequencing reads from scRNA-seq were classified as spliced or unspliced using velocyto (6). Based on this classification, RNA velocity was inferred with the stochastic model of scvelo (7) after filtering out genes with less than spliced and unspliced 20 mRNA counts in total. Diffusion components were computed with a scanpy (8) implementation of diffusion map (9). Hallmark gene signatures were retrieved from MSigDB (10) and scored using scanpy. All reported linear correlations between diffusion axis and genes or hallmarks were significant ($p < 0.05$) according to spearman correlation.

### Supplemental Figures and legends:

### Figure E1

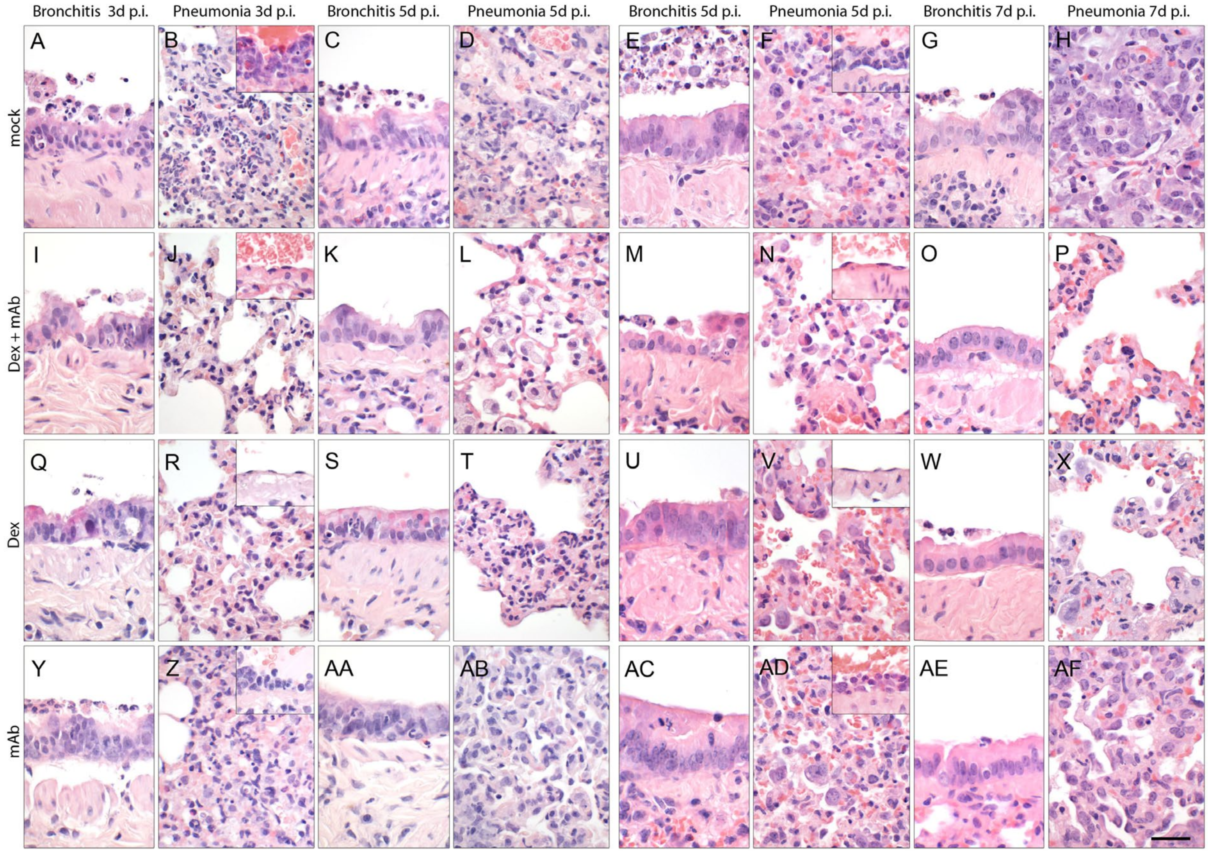


Histopathology of the left lungs of Roborovski hamsters and Syrian hamsters at 3 and 5 or 5 and 7 days, respectively, after infection with wild type SARS-CoV-2, hematoxylin and eosin stain. (A – D) Lungs of mock treated Roborovski hamsters at both time points had marked and multifocal to coalescing bronchointerstitial pneumonia. (A, C) The bronchiolar epithelium occasionally appeared irregularly hyperplastic with single cell necrosis and accumulation of cellular debris in the bronchial lumen. (B) Alveoli were infiltrated by macrophages, neutrophils and lymphocytes, in addition to necrosis of alveolar epithelial cells and hemorrhage at 3 days. Blood vessels developed marked endothelialitis predominantly at day 3 (inset). (D) At day 5, alveolar type II epithelial cells were hyperplastic. (I – L) Syrian hamsters treated with a combination of Dex + mAb developed milder lesions at both time points. (I) At 3 days after infection, bronchi had only moderate necrosis of bronchial epithelial cells as well as milder transmigration of neutrophils into the bronchial lumen whereas (J) alveoli revealed random areas of mild necrosis and infiltration by only very few macrophages and neutrophils with mild alveolar edema. Endothelialitis was not observed at all. At day 5 after infection (K) bronchial epithelium was only mildly hyperplastic (L). Activated alveolar macrophages were the dominating cells in the alveolar lumen, eliminating debris of immune cell responses. (Q - T) Treatment with Dex alone also had an advantageous affect compared to the non-treated group and resembled the histopathology patterns of the combination treated group. (Q) Bronchial changes were characterized by mild to moderate, necrotizing and suppurative bronchitis at day 3 and (S) minimal bronchitis at day 5. (R) Alveolar changes also included mild alveolar epithelial necrosis as well as moderate infiltration with macrophages, neutrophils and lymphocytes at day 3. There was no endothelialitis (inset). (T) Dex treated hamsters had increased alveolar macrophages and areas of interstitial thickening with mixed inflammatory cell infiltrates at day 5. (Y – AB) Lungs of mAb treated hamsters developed moderate to marked, extensive necrosuppurative, bronchointerstitial pneumonia with only one animal reaching the second endpoint (AA – AB) at day 5. (Y) In addition to the typical moderate and necrosuppurative bronchitis, (Z) there was moderate alveolar epithelial cell necrosis with marked infiltration by neutrophils, macrophages and lymphocytes as well as moderate to marked necrosis of alveolar epithelial cells. Blood vessel adjacent to the affected areas had moderate endothelialitis (inset). The hamster that reached the day 5 endpoint had only mild, necrosuppurative bronchitis (AA) and moderate hyperplasia of alveolar epithelial cells type II (AB).

(E – F) Lungs of mock treated Syrian hamsters at day 5 after infection had patchy areas of bronchointerstitial pneumonia and epithelial cell necrosis. (E) Bronchi showed signs of bronchial epithelial hyperplasia as well as intraluminal accumulation of cellular debris originating from deeper airways. (F) Alveoli presented with consolidation and marked interstitial as well as intra-alveolar infiltration with macrophages, lymphocytes and neutrophils, accompanied by necrosis of alveolar epithelial cells. Blood vessels adjacent to affected areas developed moderate endothelialitis. Alveolar type II epithelial cells were hyperplastic. (G – H) At day 7 after infection mock treated lungs still had multifocally consolidated areas as well as (G) bronchial epithelial hyperplasia. (H) Lungs at later time points had less immune cell infiltrates but showed severe hyperplasia of alveolar type II epithelial cells with prominent mitotic activity. (M - P) Lungs of Syrian hamsters treated with a combination of Dex + mAb developed no or only little consolidation at 5 and 7 days. (M) Sporadic single cell necrosis was detected in the bronchial epithelium at 5 days. (O) No significant changes were detected at day 7 anymore. (N) Macrophages were the predominant cell type at day 3 in the alveolar lumen, eliminating immune cell debris. Endothelialitis was absent (inset). (N, P)

Alveolar walls appeared mildly expanded at both time points. (U - X) Syrian hamsters treated with Dex developed only patchy areas of interstitial thickening but no parenchymal consolidation. (U) The bronchial epithelium appeared moderately hyperplastic a day 5 and (W) returned to normal thickness at day 7. (V, X) Alveolar walls were only mildly thickened and infiltrated by moderate numbers of neutrophils, macrophages and lymphocytes. Alveolar epithelial cells were mildly hyperplastic and endothelialitis was absent (V, inset). (AC - AF) In contrast, mAb alone treated lungs developed stronger bronchointerstitial pneumonia with areas of markedly consolidated parenchyma and (AC, AE) bronchial epithelial hyperplasia at both time points. (AD, AF) Alveolar changes were similar to lesions observed in the non-treated group, including moderate infiltration with macrophages, neutrophils and lymphocytes as well as moderate hyperplasia of alveolar type II epithelial cells and moderate endothelialitis in blood vessels adjacent to inflamed areas (AD, inset). Scale bar for all = 25 μm

### Figure E2

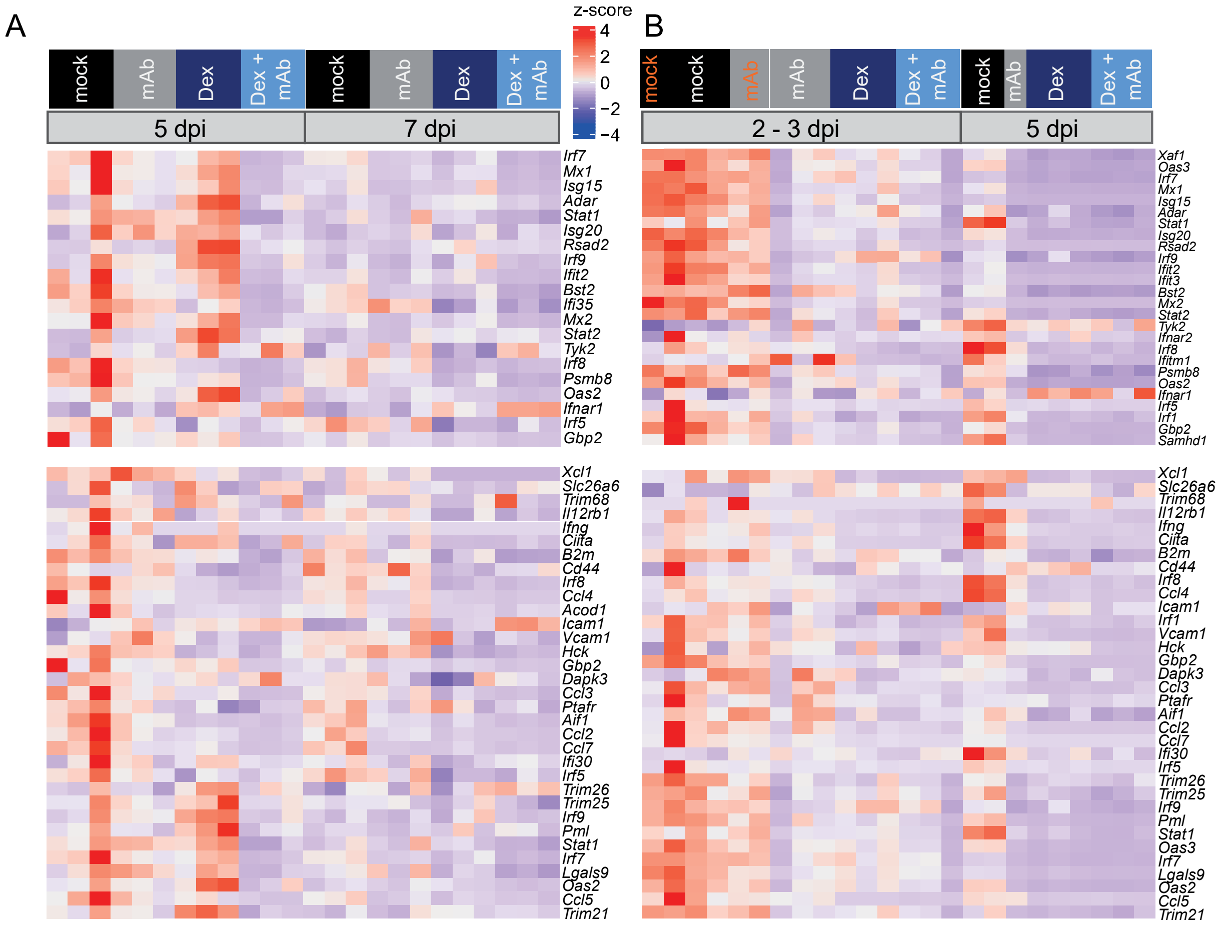


Gene expression was quantified using polyA RNA high-throughput sequencing from Syrian hamster (A) and Roborovski hamster (B) lung samples. Shown are z-scores of fpkm values calculated over all samples on a color scale ranging from blue (-4) to red (+4). Time points and treatments are shown on top of the heatmap. Samples from animals taken out at 2 dpi are shown in orange (B). The displayed genes are those from the type I Interferon / Interferon gamma response set that are annotated in the respective genomes.

**Figure E3A – E…**

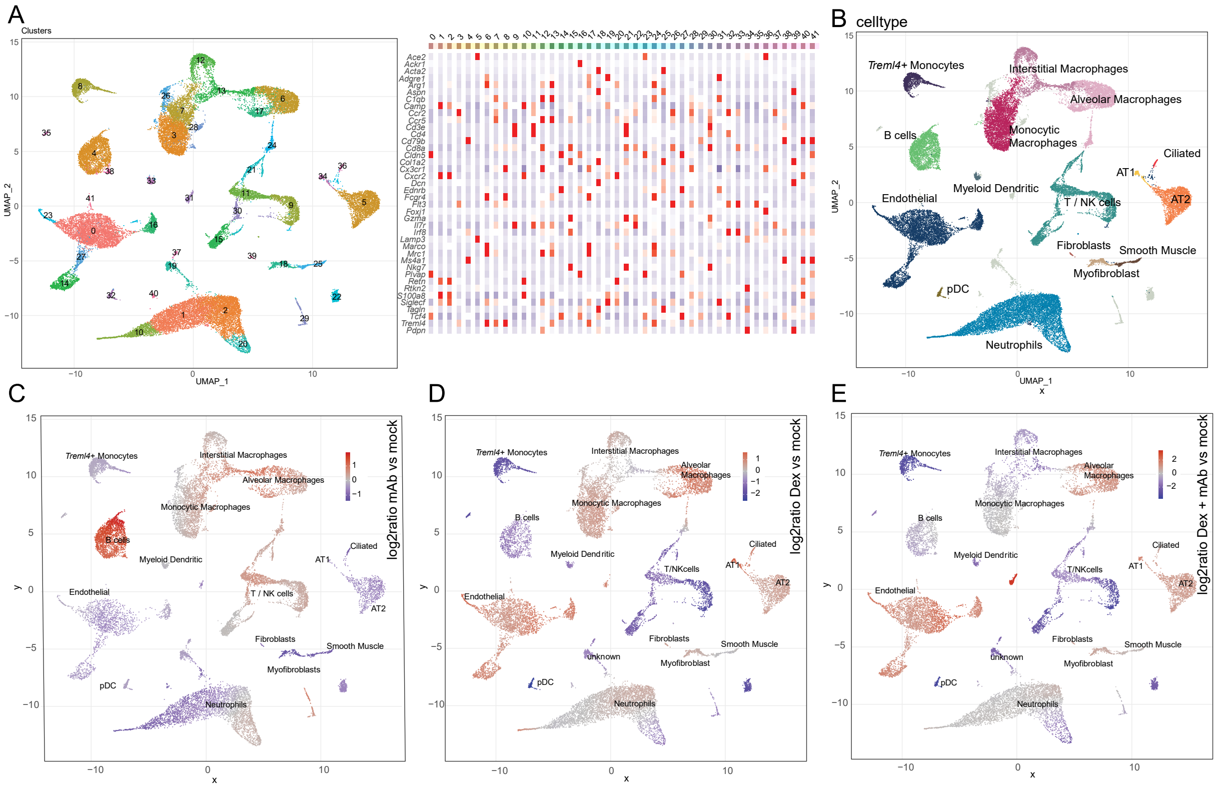


(A) Left, cells from all single-cell RNA-sequencing samples were integrated and clustered using the Louvain algorithm based on their individual transcriptomes, and two-dimensional projections performed using the UMAP algorithm. Cells were colored by their cluster identity. Right, the expression of cell type marker genes per cluster was displayed as a heatmap. Based on marker gene expression, clusters were manually assigned to cell types and were colored by cell type (B). Changes in cellular density on the UMAP projection were calculated, and cells colored by fold changes of the indicated treatment vs. mock treatment, mAb vs mock (C), Dex vs. mock (D), and Dex + mAb vs mock (E). Red indicates increased density, and blue indicates decreased density, as for example the amount of T / NK cells is reduced upon dexamethasone treatment (D). mAb: monoclonal antibody, Dex: dexamethasone, vs: versus.

**…Figure E3F – K**

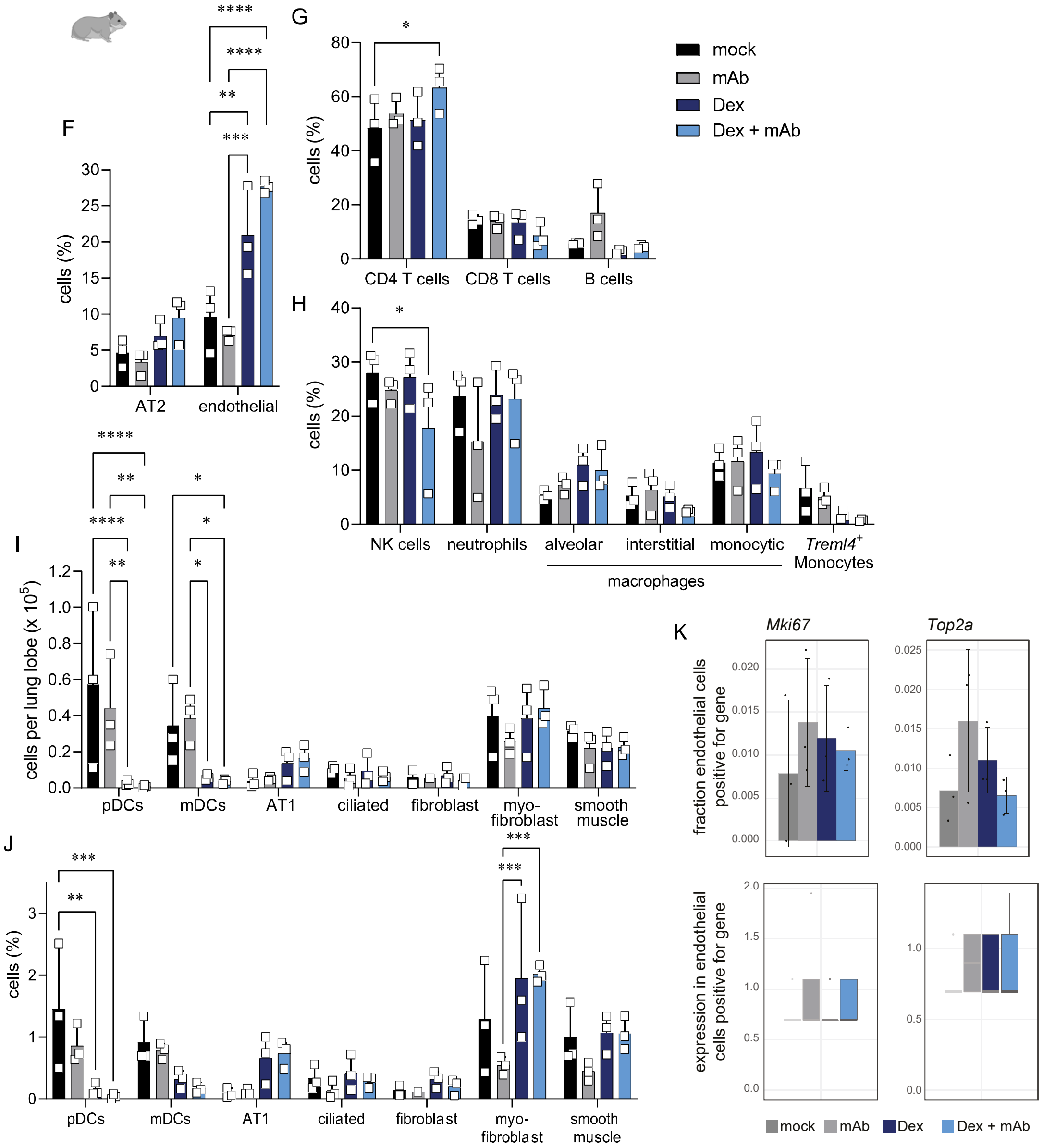


Percentage of indicated cells per lung lobe in hamsters at 2 dpi depending on treatment with mock, mAb, DEx, and Dex + mAb; AT2 and endothelial cells (F), T and B lymphocytes (G) and various innate immune cells (H). Percentages (I) and calculated numbers of indicated small cell populations. Data from scRNA-Seq and lung cell counting (Fig. 2A). Data display means ± SD. n = 3 per group. (F – J) Two-way ANOVA, Tukey's multiple comparisons test. * $P < 0.05$, ** $P < 0.01$, *** $P < 0.001$, **** $P < 0.0001$. (K) Expression of proliferation marker genes *Mki67* and *Top2a* in endothelial cells. Shown are the fraction of cells with ≥ one mRNA count (top, means ± SD, n = 3) and the expression levels in the cells with ≥ one mRNA count (bottom, boxplots, lower and upper hinges correspond to first and third quartiles. Whiskers extend to a maximum of 1.5 times the distance between first and third quartile. Outliers beyond are marked by single dots).

**Figure E4A – D…**

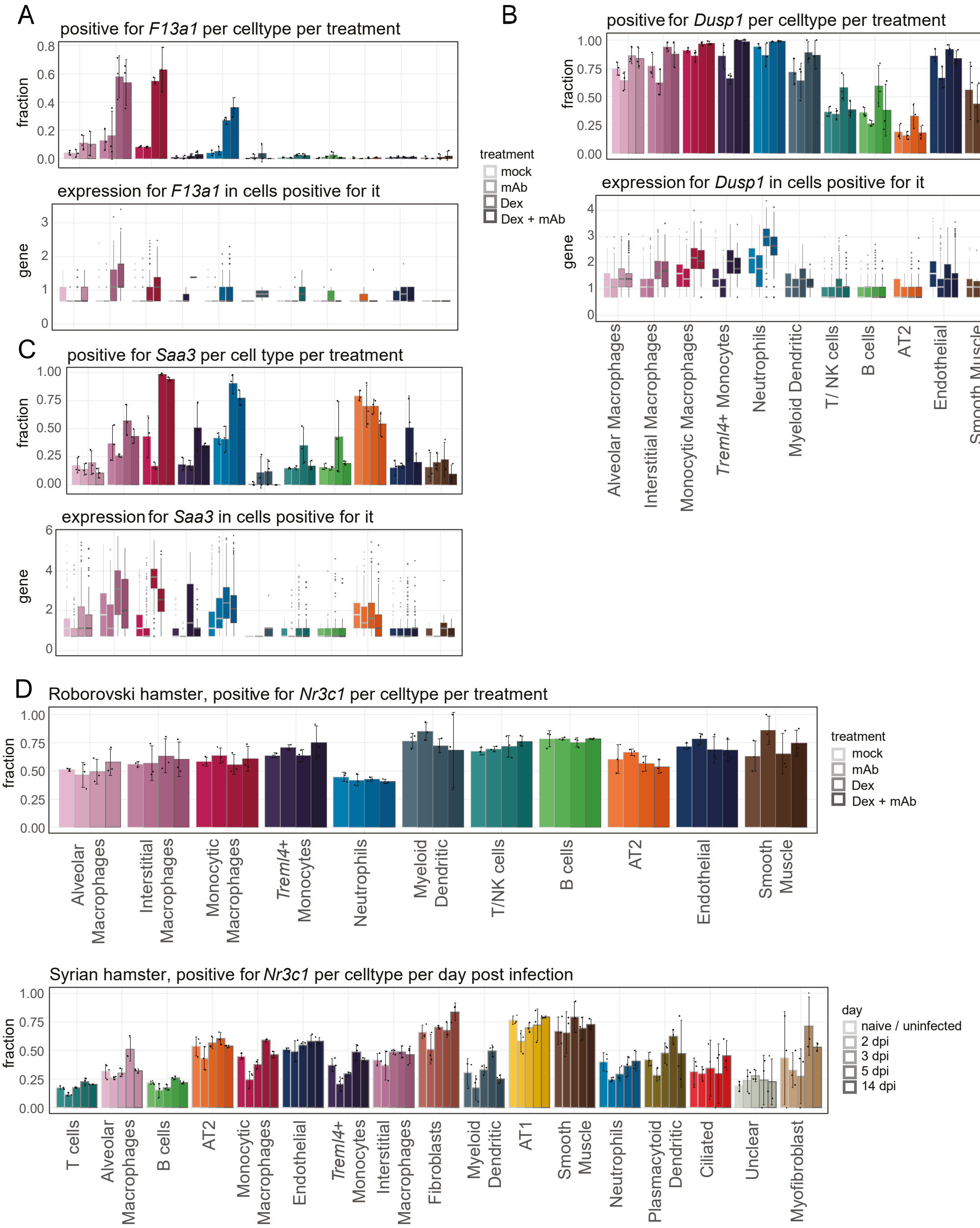


Expression of dexamethasone-induced genes *F13a1 (A)*, *Dusp1 (B)*, and *Saa3* (C) in indicated cell types. Shown are fraction of cells with ≥ one mRNA count (top panels, means ± SD, n = 3) and the expression levels in cells with ≥ one mRNA count (bottom panels, boxplots, lower and upper hinges correspond to first and third quartiles. Whiskers extend to a maximum of 1.5 times the distance between first and third quartile. Outliers beyond are marked by single dots). (D) Expression of the glucocorticoid receptor gene *Nr3c1* in the indicated cell types in Roborovski hamsters (top, this study) and Syrian hamsters (bottom, from (11)). Shown are the fraction of cells in which ≥ one mRNA count (means ± SD, n = 3).

**…Figure E4E…**

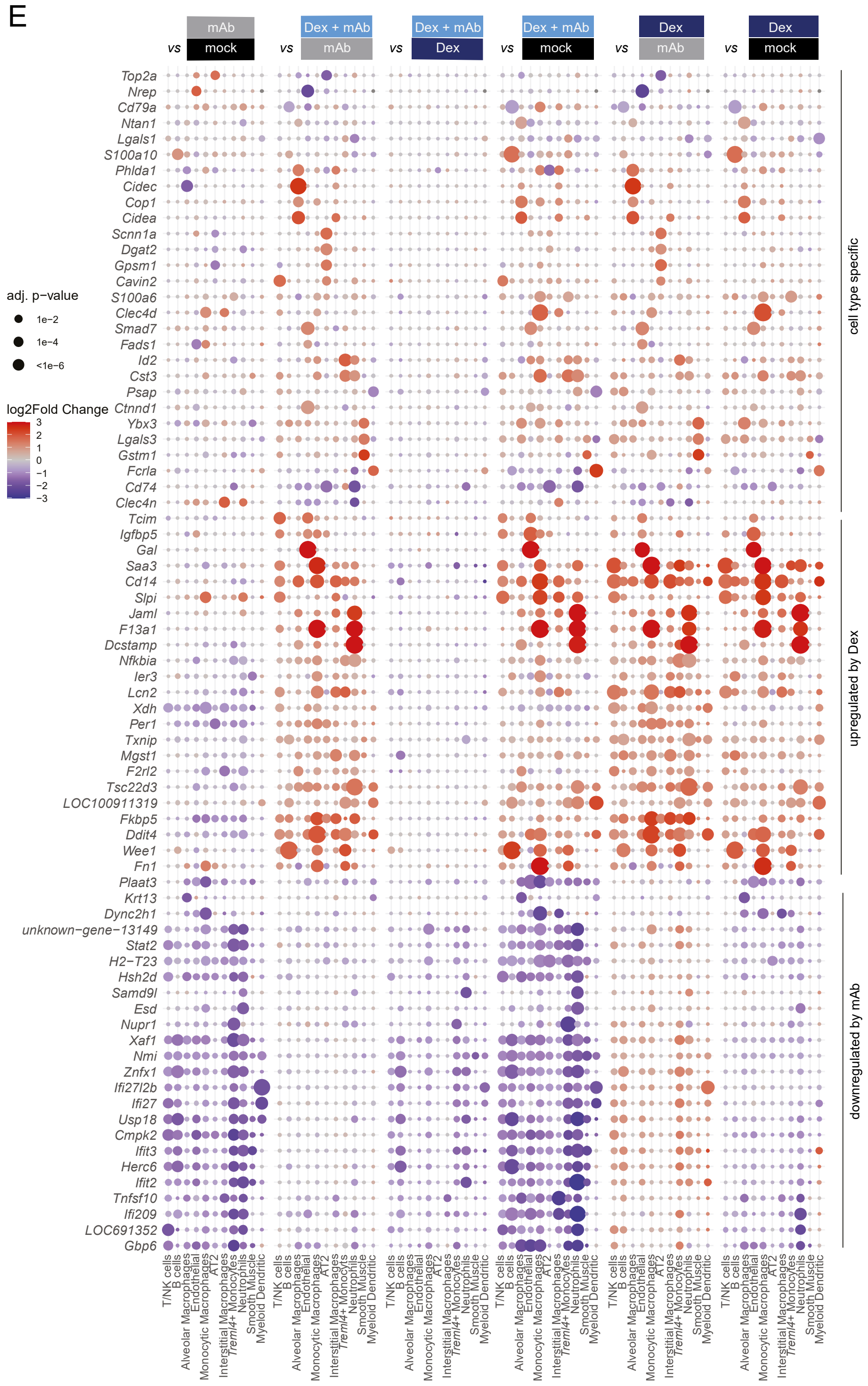


Dotplots of differentially expressed genes for all three treatments compared to each other (E). For every comparison and every cell type, the two most significantly changed genes are shown. Size and colors of the dots indicate log2-transformed fold changes (FC) and p-values, respectively. Adjusted (adj) p-values were calculated by DEseq2 using Benjamini–Hochberg corrections of two-sided Wald test p-values. Genes are ordered by unsupervised clustering.

**…Figure E4F**

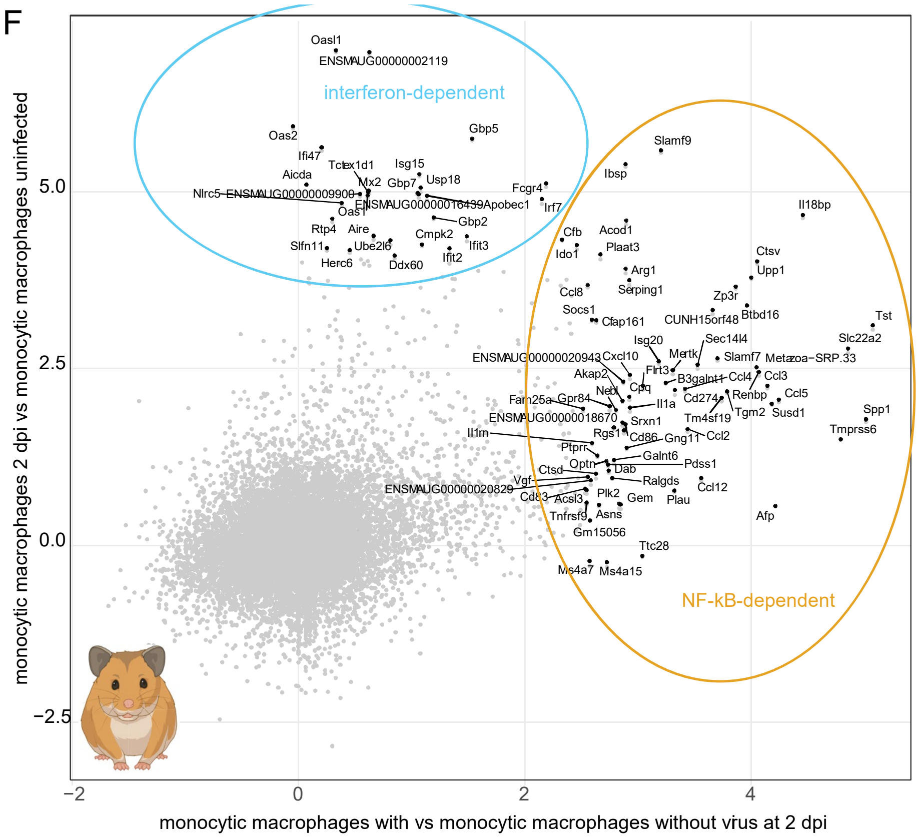


Data from: Nouailles G, Wyler E, Pennitz P, et al. Nat Commun. 2021;12(1):4869.

(F) Plotted are log2-transformed gene expression fold changes in monocytic macrophages from Syrian hamsters (from (11)) against each other, comparing 2 dpi to uninfected (vertical axis) and comparing cells containing viral RNA vs. cells without viral RNA (horizontal axis). Gene sets were defined as either predominantly NF-kB-dependent (orange oval, right) or predominantly interferon-dependent (light blue oval, top).

### Figure E5A – C…

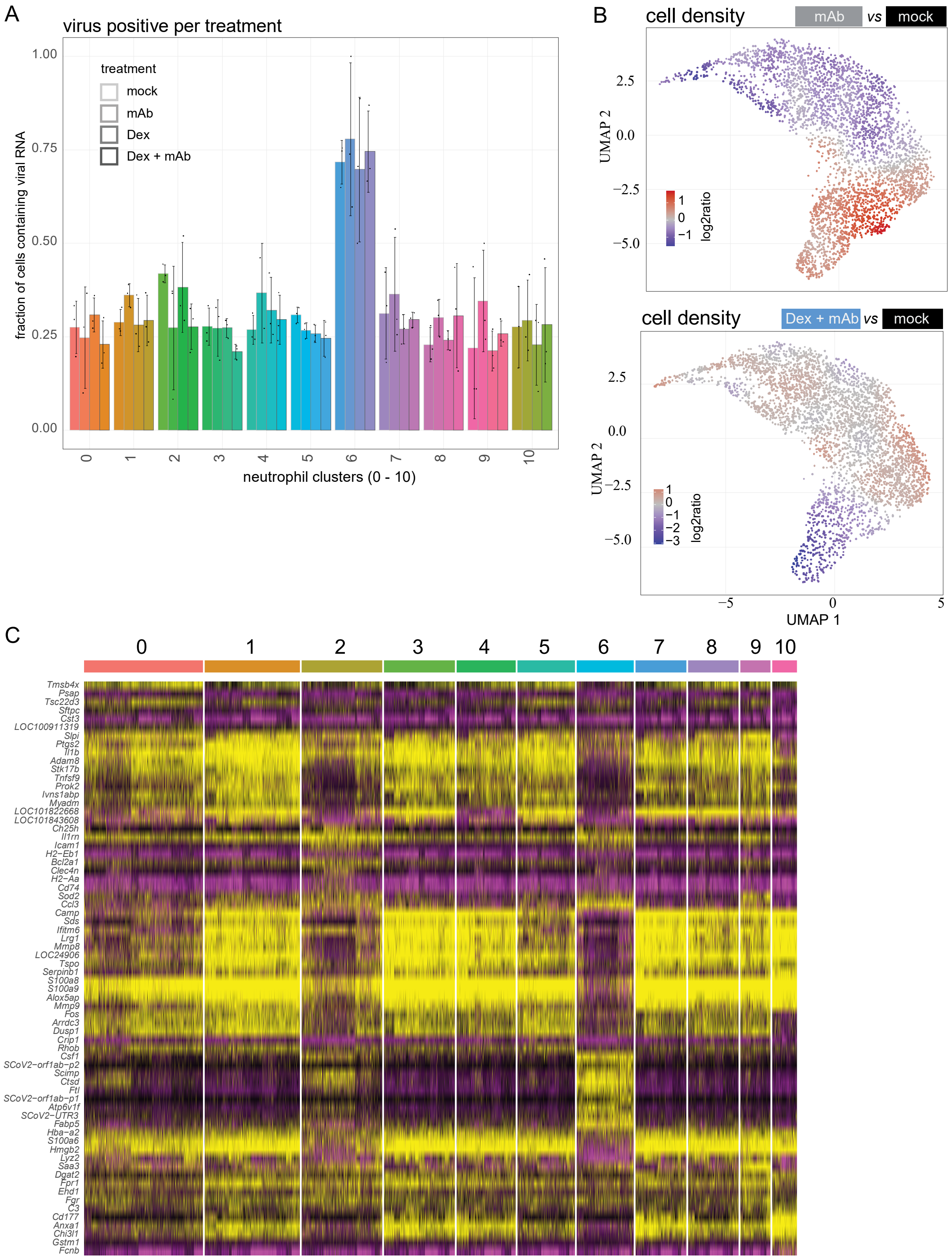


Fraction of cells in the indicated neutrophil subcluster (1 – 10) containing with ≥ one count viral RNA count, means ± SD, n = 3 (A). (B) As in Figure 4C, but for the comparisons mAb vs. mock and dex + mAb vs. mock. (C) Heatmap of the most enriched genes per indicated neutrophil subcluster (1 – 10).

### …Figure E5D – E…

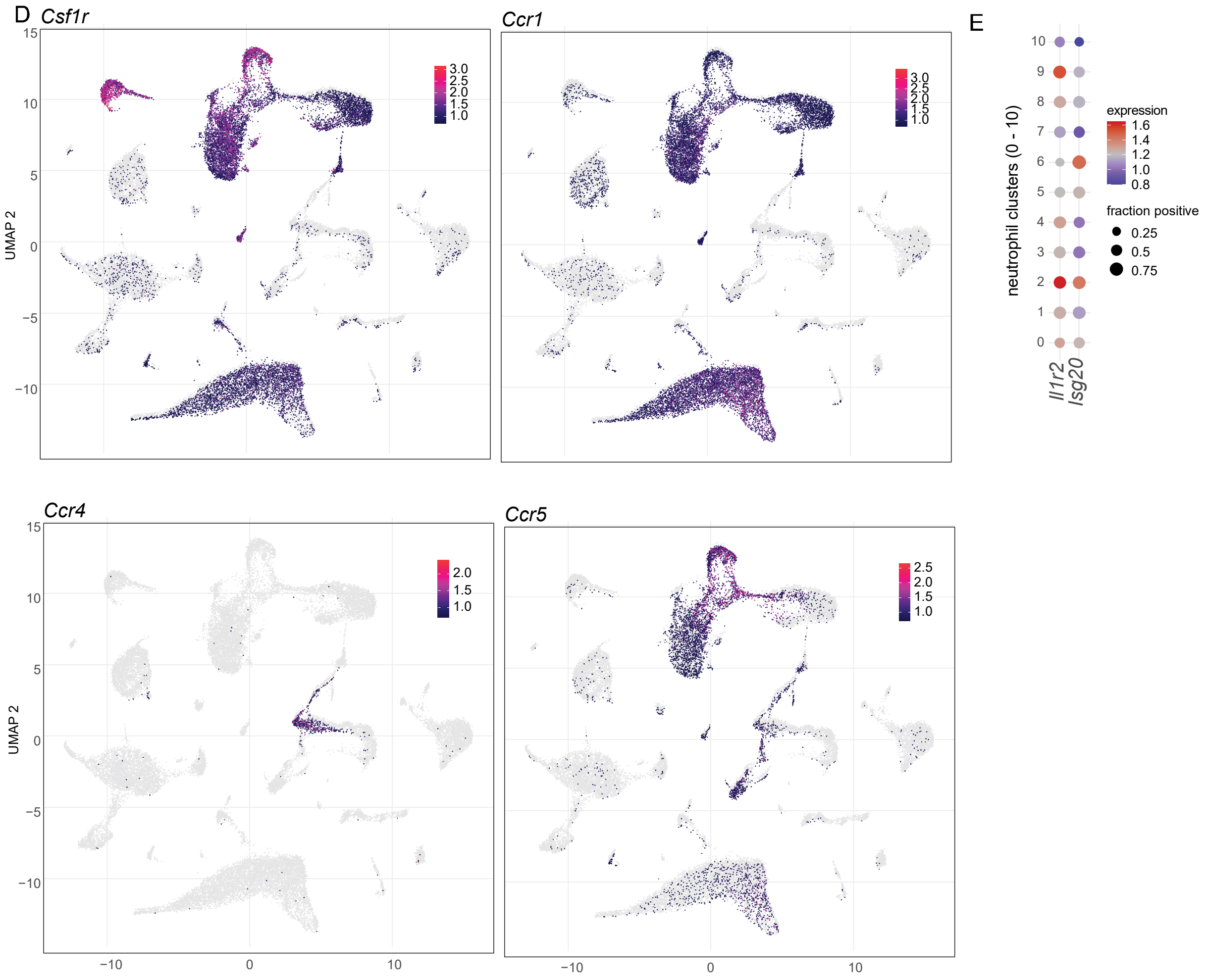


(D) Individual cells on the two-dimensional projections introduced in Figure E3A are colored by the expression values of the respective genes. (E) Expression of the indicated genes over all animals in the clusters as defined in Figure 5A. The dot size indicates the fraction of cells in the clusters as indicated on the left from mock-treated animals, with ≥ one mRNA count detected for the respective gene. The color represents the average expression in those cells.

### …Figure E5F

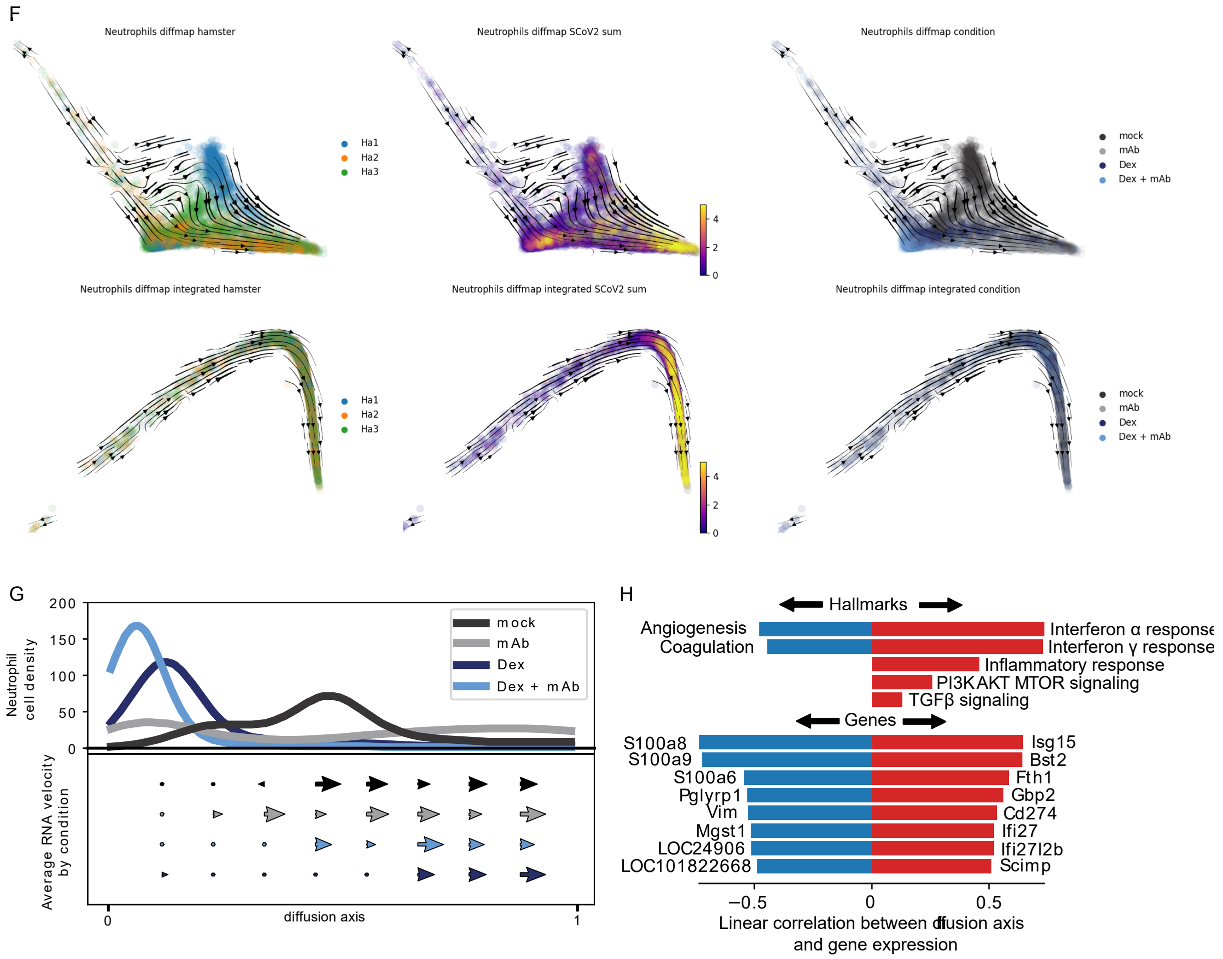


(F) Diffusion map embedding based on a principal components without (top row) or with (bottom row) integration by Seurat. The cells are colored as following. Left, by animal (note that every represents four animals, namely one of mock, one of mAb, one of Dex, one of mAb + Dex), indicating inter-individual variability on the vertical axis (top left), which is removed by the integration (bottom left). Middle, by amount of viral RNA. Right, by condition. G) Top part, kernel estimated cell density along the primary diffusion component of the non-integrated diffusion map. Bottom part, average RNA velocity per condition projected onto the diffusion component 1. (H) Hallmark signatures (10) and gene expression profiles were linearly correlated with the diffusion axis. Plotted are the correlation coefficients in either red (positive negative) or blue (negative correlation).
